# Male pheromone composition depends on larval but not adult diet in *Heliconius melpomene*

**DOI:** 10.1101/341602

**Authors:** Kathy Darragh, Kelsey J.R.P. Byers, Richard M. Merrill, W. Owen McMillan, Stefan Schulz, Chris D. Jiggins

**Affiliations:** Department of Zoology, University of Cambridge, Cambridge, United Kingdom; Smithsonian Tropical Research Institute, Panama; Division of Evolutionary Biology, Ludwig-Maximilians-Universität, Germany; Institute of Organic Chemistry, Technische Universität Braunschweig, Braunschweig, Germany

## Abstract

Condition-dependent traits can act as honest signals of mate quality, with fitter individuals able to display preferred phenotypes. Nutrition is known to be an important determinant of individual condition, with diet known to affect many secondary sexual traits. In *Heliconius* butterflies, male chemical signalling plays an important role in female mate choice. *Heliconius* pheromones are sexually dimorphic, found only in mature males, but it is unclear what information they convey to the female. Here, we manipulate both the larval and adult diet of male *Heliconius melpomene rosina* to test for environmental effects on wing and genital pheromone production. We find no evidence that adult pollen consumption affects pheromone production in the first ten days after eclosion. We also find strong overlap in the chemical profiles of individuals reared on different larval host plants. However, individual compounds were found in different amounts between host plant treatments. Further electrophysiological and behavioural experiments will be needed to determine the biological significance of these differences.

## Introduction

Sexual ornaments often act as an indicator of male quality and evolve in response to sexual selection imposed by female preferences (Zahavi, 1975, 1977; Andersson, 1986). Male “quality” can reflect both direct and indirect benefits gained by the female (Andersson, 1994). Direct benefits might include resources that increase female lifetime reproductive success, such as food, shelter, parental care or protection from predators. Indirect benefits, on the other hand, are those that increase genetic quality of a female’s offspring; here, impressive sexual ornaments reflect the ability of males to provide genes that increase the survivorship or mating success of offspring (Andersson, 1986, 1994). Females exhibiting a preference for males with the sexual ornament thus benefit from mating with a high-quality male, which in turn selects for females with this preference. Ornaments may be an honest signal of quality if they are condition-dependent, where only the best quality males are able to display the phenotype.

Many secondary sexual traits, such as the exaggerated eye-span of stalk-eyed flies (Cotton *et al.*, 2004), rhinoceros beetle horns (Emlen *et al.*, 2012), or male drumming rate of wolf spiders (Kotiaho, 2000) exhibit condition dependence. Whilst most studies have focused on visual and acoustic traits, chemical signalling has been less well studied. Diet manipulation experiments show that the overall nutritional condition of an individual is important for pheromone production in some groups, with nutrient stress affecting male pheromones in *Tenebrio* beetles (Rantala *et al.*, 2003), cockroaches (Clark *et al.*, 1997) and burying beetles (Chemnitz *et al*., 2015). Furthermore, pheromone signalling is costly, (Johansson *et al*., 2005; Harari *et al*., 2011), and diet-mediated changes can enforce signal reliability, making male pheromones a good candidate as an honest signal (Henneken *et al*., 2017).

As well as overall diet quality, which affects condition of the signaller, specific diet components can be important. A classic example of this is the consumption of carotenoids and their effect on male coloration (Endler, 1983; Hill, 1992; Hill *et al*., 2002; Shawkey *et al*., 2006). In chemical signalling, compounds sequestered in the diet can act as sex pheromones, or as pheromone precursors which the animal then metabolises into the final pheromone product (Landolt & Phillips, 1997). One well-studied example of this is the sequestration of pyrrolizidine alkaloids (PAs) from larval host plants. Males of the moth *Utethesia ornatrix* sequester PAs which are then used to produce the sex pheromone hydroxydanaidal (Conner *et al*., 1981; Eisner & Meinwald, 2003). Some PAs are transferred to the female during mating and chemically protect the eggs, providing a direct benefit (Dussourd *et al*., 1988, 1991a). The amount of male pheromone produced indicates the amount of stored alkaloid, providing an honest signal to females of the amount of egg protection she will receive (Eisner & Meinwald, 1995; Iyengar *et al*., 2001). In many cases it is probable that both overall nutrient condition and the consumption of specific compounds are both important, such as in the oriental fruit fly, where both overall protein intake and the intake of a specific precursor, methyl eugenol, affect mating success (Shelly *et al*., 2007).

Chemical cues influence the mate choice decisions of the butterfly *Heliconius melpomene rosina* Boisduval (Nymphalidae), a subspecies of *Heliconius melpomene* found in central Panama. Androconial sex pheromones have been described in the wing overlap region of sexually mature males and are sexually dimorphic (Darragh *et al*., 2017). They consist of a bouquet of compounds, including octadecanal as a main component (Mérot *et al*., 2015; Darragh *et al*., 2017; Mann *et al*., 2017). These chemical cues are important for mating, with females strongly discriminating against males which have their androconia experimentally blocked (Darragh *et al*., 2017). However, it is still unclear what information (*e*.*g*. age or male quality) is being conveyed to females by these cues.

As well as androconial pheromones, males store compounds in genital scent glands (Gilbert, 1976; Schulz *et al*., 2007, 2008; Estrada *et al*., 2011). These are transferred to the female during mating and act as anti-aphrodisiacs, repelling males to delay re-mating (Gilbert, 1976; Schulz *et al*., 2008). In *H. melpomene*, (*E*)-β-ocimene acts as an anti-aphrodisiac, with a bouquet of esters and alcohols thought to moderate evaporation rate (Schulz *et al*., 2008). (*E*)-β-ocimene is also found in large amounts in the flower and vegetation of *Lantana camara*, a tropical flowering plant often visited by *Heliconius* (Andersson *et al*., 2002), and elicits strong antennal responses in females (Andersson & Dobson, 2003). Reduced harassment by other males is thought to be beneficial to the female following mating, and so these compounds could lead to a direct benefit to females by advertising the female’s mated status. Longer-term, there may be conflict over when a female should re-mate (Andersson *et al*., 2000, 2004). This leads to sexual conflict within the system over the timing of re-mating, as supported by rapid evolution of genital pheromone composition (Estrada *et al*., 2011). Despite this clear role of genital compounds in male deterrence, the role of these same compounds in female choice remains unclear. Females may benefit from choosing males that have a lower amount of (*E*)-β-ocimene, allowing them to re-mate again sooner. Whilst the dynamics of the costs and benefits of this are unclear, it is quite likely that the genital compounds are involved in female choice.

To test for condition dependent effects, we need to manipulate environmental factors. Although we do not know exactly when the androconial or genital compounds are produced, they are not present in freshly-eclosed males (Schulz *et al*., 2008; Darragh *et al*., 2017), and so both larval and adult diet could be important for their production. Feeding experiments with chemically labelled precursors showed that (*E*)-β-ocimene can be synthesized by adult H. melpomene (Schulz *et al*., 2008). In Panama, H. melpomene rosina is a host plant specialist, and females oviposit almost exclusively on Passiflora menispermifolia (Merrill *et al*., 2013). Host plant use could affect the pheromone bouquet if larval sequestration of specific compounds, or compound precursors, from the host plant is necessary. Heliconius raised on their preferred host plant may have a higher quality diet (Smiley, 1978), and so pheromone production could also be increased due to higher overall quality of the individual. Uniquely among butterflies, adult Heliconius are able to feed on pollen, providing a source of amino acids (Gilbert, 1972), which might also be necessary for pheromone production.

Here, we investigated how larval and adult diet affect the chemical profile of male *H. melpomene rosina*, a monophagous population from central Panama. We reared larvae on three different *Passiflora* species. *P. menispermifolia*, the preferred host plant, *P. vitifolia*, and *P. platyloba*. The latter two species are not used by *H. melpomene* in the wild in Panama but are potential hosts and larvae survive well on both. *P. vitifolia* is common in the Gamboa area, while *P. platyloba* was collected from the Darien region of Panama. In a second experiment, we maintained adult male *H. melpomene* with and without access to pollen. In both experiments, we analysed chemical extracts from both the androconial and genital regions of sexually mature male butterflies.

## Methods

*Heliconius melpomene rosina* were reared at the Smithsonian Tropical Research Institute (STRI) facilities in Gamboa, Panama. Outbred stocks were established from wild individuals collected in Gamboa (9°7.4’ N, 79°42.2’ W, elevation 60 m) in the nearby Soberania National Park and in San Lorenzo National Park (9°17’N, 79°58’W; elevation 130 m).

For the host plant experiment, larvae were reared on either *Passiflora platyloba*, *P. vitifolia* or *P*. *menispermifolia.* Adult butterflies were kept in cages with other males and were provided with a ~20% sugar solution containing bee pollen and with *Psychotria poeppigiana*, *Gurania eriantha*, *Psiguiria triphylla*, and *Psiguria warscewiczii* as pollen sources. For the pollen experiment, all larvae were reared on *Passiflora platyloba*. Adults were then randomly divided into two groups. The first was provided with a ~20% sugar solution and the above plant species as pollen sources; the second group was only provided with a ~20% sugar solution and no pollen source.

Wing and genital tissue of mature male individuals (10-12 days post-eclosion) was collected between February 2016 and April 2017 for chemical analysis. The individuals raised on *Passiflora platyloba* and provided with pollen are previously published samples (Darragh *et al.*, 2017). To account for a potential difference in growth rate of individuals reared on different host plants, we measured the forewing length of adult butterflies for the host plant experiments. The hindwing androconial region of the wing, previously described as the grey-brown overlapping region of the wing (Darragh *et al.*, 2017), was dissected for analysis. To extract pheromones, the tissue was soaked in 200µl dichloromethane containing 200ng 2-tetradecyl acetate in 2ml glass vials with PTFE-coated caps (Agilent, Santa Clara, USA) for one hour. The solvent was then transferred to new vials and stored at −20°C. Samples were evaporated under ambient conditions at room temperature prior to analysis. Pheromone extracts were analysed by GC/MS using an Agilent model 5977 mass-selective detector connected to a Agilent GC model 7890B and equipped with a Agilent ALS 7693 autosampler. HP-5MS fused silica capillary columns (Agilent, Santa Clara, USA, 30 m × 0.25 mm, 0.25 μm) were used. Injection was performed in splitless mode (250°C injector temperature) with helium as the carrier gas (constant flow of 1.2 ml/min). The temperature programme started at 50°C, was held for 5 minutes, and then rose at a rate of 5°C/minute to 320°C, before being held at 320°C for 5 minutes. Components were identified by comparison of mass spectra and gas chromatographic retention index with those of authentic reference samples and also by analysis of mass spectra. Components were quantified using 2-tetradecyl acetate as an internal standard. As in previous analysis, only compounds eluting earlier than hexacosane were analysed in wing samples (Darragh *et al.*, 2017). Later compounds were identified as cuticular hydrocarbons, 2,5-dialkyltetrahydrofurans, cholesterol and artefacts (*e.g.* phthalates or adipates). The variability in the late eluting cuticular hydrocarbons was low and did not show characteristic differences between samples.

The samples were visualised using a non-metric multidimensional scaling (NMDS ordination, based on a Bray-Curtis similarity matrix, in three dimensions. This was carried out using the metaMDS function in the R-package vegan version 2.5-1 (Oksanen *et al*., 2017), with visualisation using the R-package ade4 (Dray & Dufour, 2007). We tested for homogeneity of dispersion (a measure of the variation, or spread, of the group) between groups using the betadisper and permutest functions to perform a permutation multivariate ANOVA. To compare overall chemical composition between groups, we carried out PERMANOVA (permutational multivariate analysis of variance) testing. This was also performed using a Bray-Curtis similarity matrix, with 1000 permutations, using the adonis2 function in the package vegan (Oksanen *et al*., 2017). The “margin” option in adonis2 was used to determine the effect of each term in the model, including all other variables, to avoid sequential effects. To test for differences between groups for individual compounds we used non-parametric Kruskal-Wallis tests. To correct for multiple-testing, we used the p.adjust function in R, with false detection rate (fdr) correction, which controls for the proportion of false positives. All statistical analyses were performed with *R* version 3.3.1 (R Core Team, 2016).

## Results

In total, we sampled the androconia of 96, and the genitals of 91, mature adult butterflies. 43 androconial and 44 genital samples were collected from adult butterflies raised from larvae reared on different host plants. 53 androconial and 47 genital samples were collected from adult butterflies kept with or without access to pollen. The minimum group size for any treatment was 11 individuals.

For the individuals reared on different host plants we also measured their forewing length at the time of sampling to test for effects of diet on size. One individual was excluded from this analysis as the wings were torn. We found that larval host plant affects forewing length (ANOVA, df=2, F=3.755, p=0.032, Figure S1), with adults which were reared as larvae on *P. menispermifolia* having a mean forewing length of 3.54cm, those on *P. platyloba* had a mean of 3.52cm and on *P. vitifolia* a mean of 3.36cm. However, post-hoc Tukey comparisons did not find any pairwise significant difference between groups.

### Chemical analysis

We initially analysed the androconia of 20, and genitals of 18, *H. melpomene* reared on *P. platyloba*. The most abundant compounds found in the androconia are syringaldehyde, octadecanal, octadecan-1-ol, (*Z*)-11-icosenal, and icosanal, as found previously (Fig. S2, Table S1)(Darragh *et al*., 2017). These compounds are present with a mean of greater than 100ng per individual, with octadecanal found in the highest amounts (mean 740ng).

The genital region is dominated by one main compound, (*E*)-β-ocimene, which is found in amounts 20 times greater than any other genital compound (mean 37.389µg), as previously reported (Schulz *et al.*, 2008). It is found alongside a bouquet of other terpenes, alcohols, aromatic compounds, macrolides, esters and alkanes (Fig. S3, Table S2)(Schulz *et al.*, 2008).

There is little overlap in compounds found between the two regions, with only ten out of 117 compounds found in both. The genital region contains higher amounts of compounds, and overall more compounds, with 80 in the genitals compared to 47 in the androconia. The most abundant genital compound, (*E*)-β-ocimene, is more volatile (has a higher vapour pressure) than the main compounds found in the androconial region.

### Host plant experiments

Our experiments revealed that *H. melpomene* reared on *P. platyloba* (20 individuals), *P. menispermifolia* (11 individuals), or *P. vitifolia* (12 individuals) did not differ significantly in their overall androconial pheromone bouquet (PERMANOVA, df=2, F=1.791, p=0.080, Figure 1A, Table S1), and no significant difference in dispersion between groups was detected (ANOVAnce, df=2, F=1.384, p=0.262). However, when we look at the individual compounds in each treatment, over one quarter (12/47), are present in significantly different amounts between groups (Table 1).

**Figure 1.**
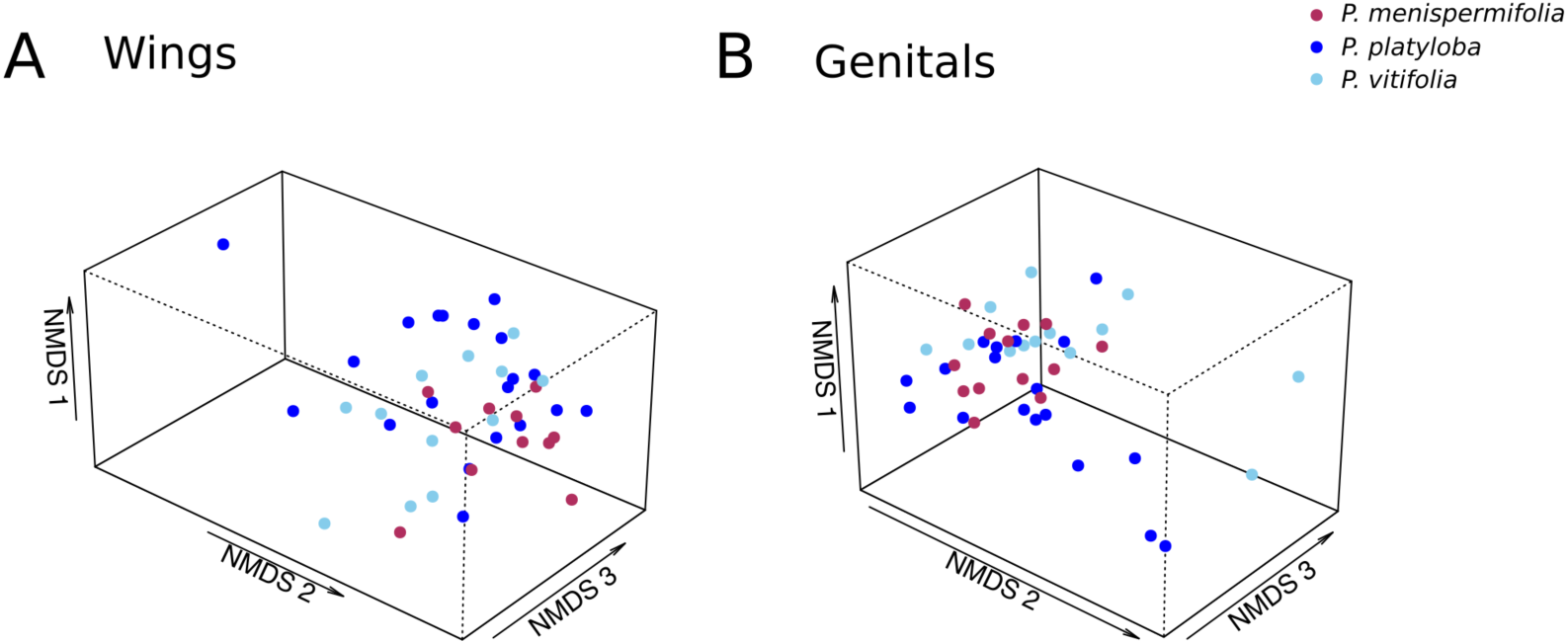
NMDS (non-metric multidimensional scaling) plot illustrating in three dimensions the overlapping variation in chemical compounds of male *H. melpomene* raised on three different *Passiflora* species. (A) Wing compound bouquets do not differ significantly after 10 days (PERMANOVA, n = 43, df=2, F=2.429, p=0.023). Stress=0.140. (B) Genital compound bouquets do not differ significantly after 10 days (PERMANOVA, n = 44, df=2, F=1.184, p=0.318). Stress=0.099.

**Table 1:**
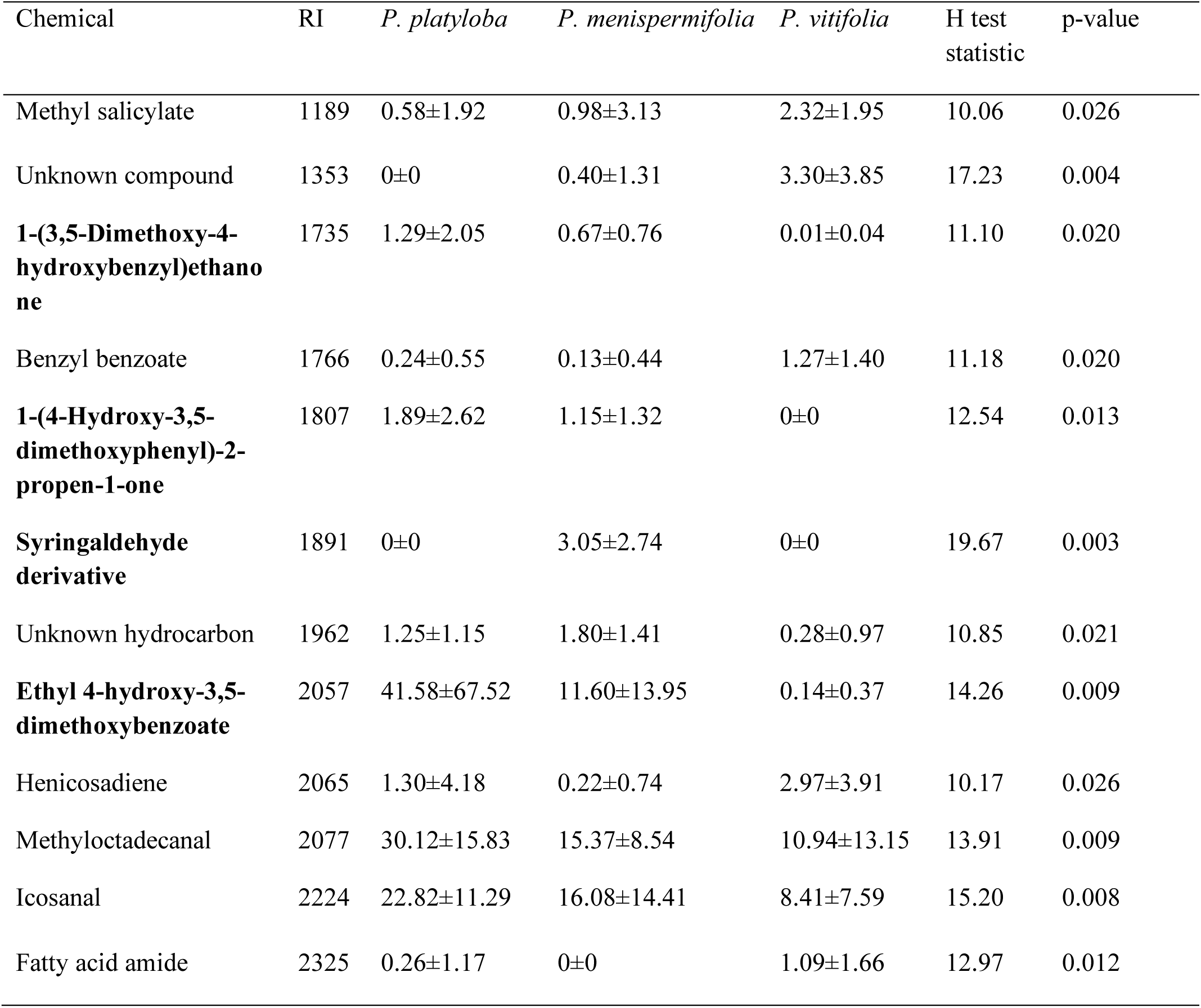
Androconial compounds which significantly differed between *H. melpomene* reared on different host plants. The gas chromatographic retention index (RI) is reported for each compound. Mean amounts (ng) ± standard deviation, as well as Kruskal-Wallis non-parametric test p-values (false detection rate corrected) are provided. Compounds in bold are predicted to be plant-derived.

Genital compounds of *H. melpomene* reared on *P. platyloba* (18 individuals), *P. menispermifolia* (13 individuals), or *P. vitifolia* (13 individuals) were also not found to differ overall between host plant treatments (PERMANOVA, df=2, F=1.184, p=0.318, Figure 1B, Table S2). The dispersion of individuals between treatments did differ (PERMANOVA, df=2, F=3.577, p=0.037), with pairwise permutation analysis revealing that the dispersion of individuals raised on *P. menispermifolia* is different from both *P. vitifolia* and *P. platyloba*, which do not differ from each other (Table S3). NMDS visualisation reveals less variation between individuals reared on the monophagous host *P. menispermifolia* (Figure 1B). Furthermore, seven out of 80 individual compounds differ between groups (Table 2).

**Table 2:**
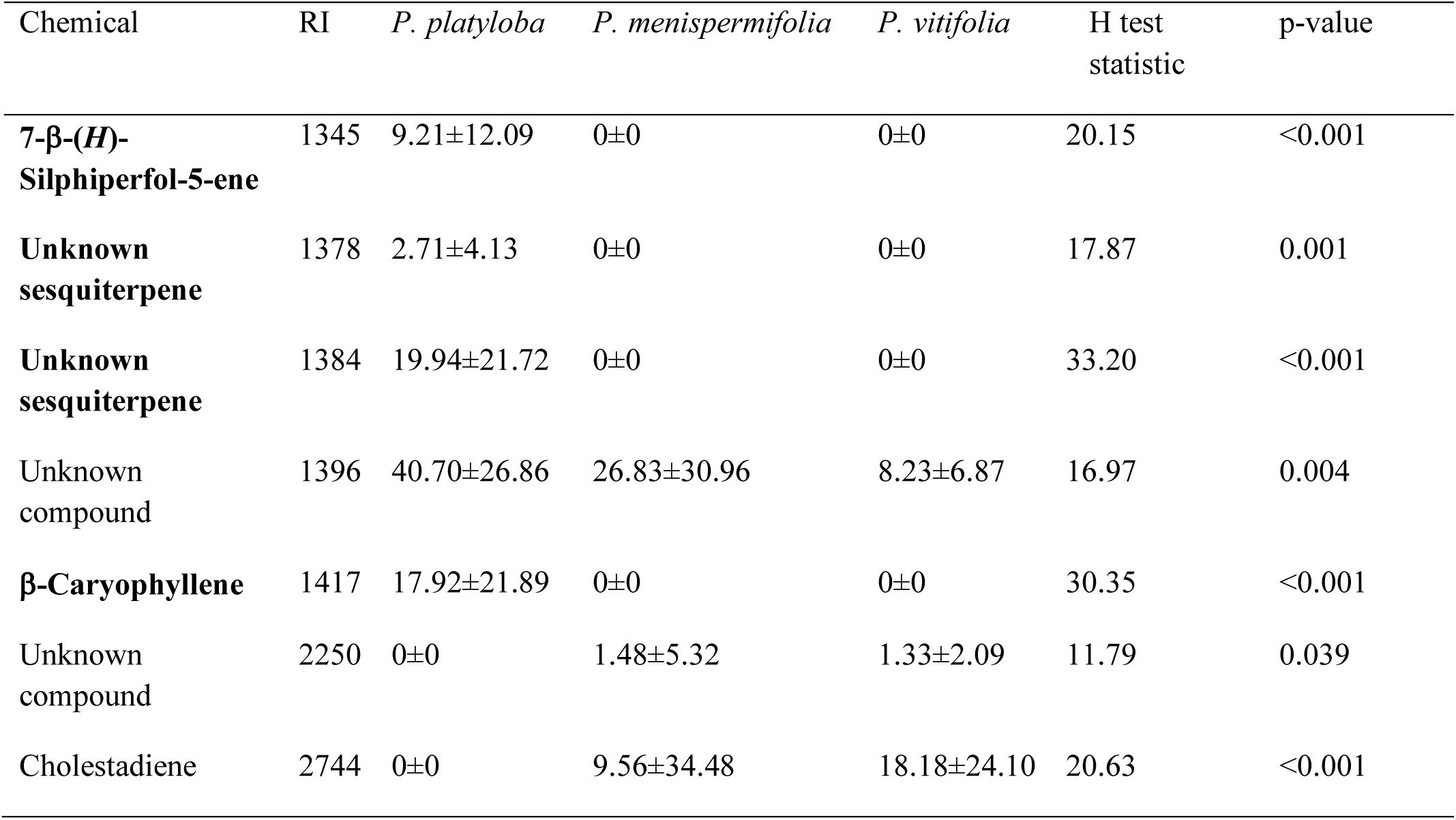
Genital compounds which differed significantly between *H. melpomene* reared on different host plants. The gas chromatographic retention index (RI) is reported for each compound. Mean amounts (ng) ± standard deviation, as well as Kruskal-Wallis non-parametric test p-values (false detection rate corrected) are provided. Compounds in bold are predicted to be plant-derived.

| Chemical | RI | *P. platyloba* | *P. menispermifolia* | *P. vitifolia* | H test statistic | p-value |
| --- | --- | --- | --- | --- | --- | --- |
| **7-β-(H)-Silphiperfol-5-ene** | 1345 | 9.21±12.09 | 0±0 | 0±0 | 20.15 | <0.001 |
| **Unknown sesquiterpene** | 1378 | 2.71±4.13 | 0±0 | 0±0 | 17.87 | 0.001 |
| **Unknown sesquiterpene** | 1384 | 19.94±21.72 | 0±0 | 0±0 | 33.20 | <0.001 |
| Unknown compound | 1396 | 40.70±26.86 | 26.83±30.96 | 8.23±6.87 | 16.97 | 0.004 |
| **β-Caryophyllene** | 1417 | 17.92±21.89 | 0±0 | 0±0 | 30.35 | <0.001 |
| Unknown compound | 2250 | 0±0 | 1.48±5.32 | 1.33±2.09 | 11.79 | 0.039 |
| Cholestadiene | 2744 | 0±0 | 9.56±34.48 | 18.18±24.10 | 20.63 | <0.001 |

To account for the effect of host plant on size we incorporated forewing length into the PERMANOVA model. Wing size was not a significant factor influencing chemical composition of *H. melpomene* genitals (PERMANVO, df=1, F=0.357, p=0.814), but did influence androconial chemical bouquets (PERMANOVA, df=1, F=4.234, p=0.009), accounting for approximately 9% of the variation.

### Pollen experiments

*H. melpomene* butterflies reared with or without pollen for 10 days do not differ in either androconial (PERMANOVA, df=1, F=1.653, p=0.160), or genital (PERMANOVA, df=1, F=1.259, p=0.102) whole chemical bouquets (Figure 2, Table S4, Table S5). We analysed the androconia of 20 individuals reared with pollen and 33 without, and also the genitals of 20 individuals reared with pollen, and 27 without. False detection rate corrected Kruskal-Wallis testing found no compounds in significantly different amounts between the groups.

**Figure 2.**
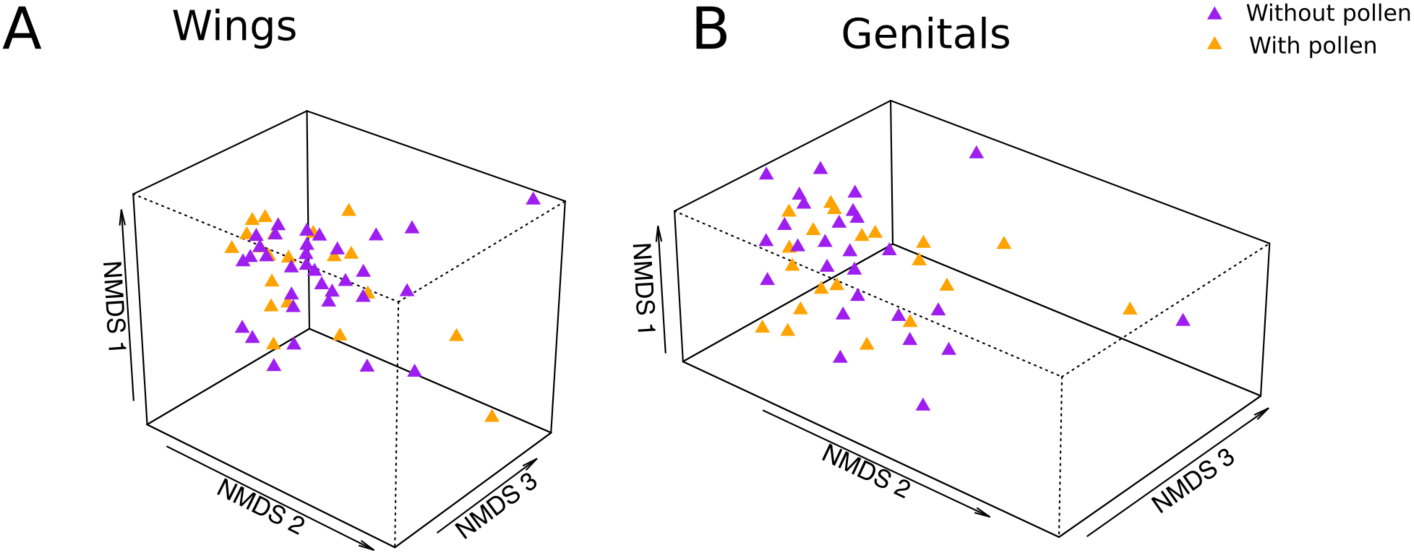
NMDS (non-metric multidimensional scaling) plot illustrating in three dimensions the overlapping variation in chemical compounds of male *H. melpomene* raised with or without pollen. (A) Wing compound bouquets do not differ significantly after 10 days (PERMANOVA, n =53, df=1, F=1.653, p=0.160). Stress=0.131. (B) Genital compound bouquets do not differ significantly after 10 days (PERMANOVA, n = 47, df=1, F=1.259, p=0.102). Stress=0.108

## Discussion

Chemical signalling is known to be important for intraspecific female mate choice in *Heliconius* (Darragh *et al*., 2017). The information conveyed by these signals, such as age, species identity, or mate quality, remains unclear. In this study, we test the effect of both larval and adult diet on male pheromone composition. We find no evidence that adult pollen consumption affects male pheromone production in the first ten days of adult life. In contrast, individual wing and genital compounds were found in different amounts between larval host plant treatment groups, and dispersion varied between host plant treatments for genitals.

Our results broadly confirm previous identification of compounds produced by *H. melpomene*. The most abundant compounds identified in *H. melpomene* androconia and genitals are the same as those identified by previously published studies (Schulz *et al*., 2008; Estrada *et al*., 2011; Mérot *et al*., 2015; Darragh *et al*., 2017; Mann *et al*., 2017). We did not identify all the compounds found previously in genitals (Schulz *et al*., 2008). This is likely due to variation in sample collection, as the previous compound list was derived from pooled samples, allowing for a higher detection threshold. In the androconia, we did not detect ethyl palmitate, ethyl oleate or ethyl stearate, previously reported compounds (Mann *et al*., 2017). We did detect ethyl oleate in the genitals suggesting that previous reports in the androconia were due to contamination from genital contact. We also found many more compounds, probably due to improved GC-MS detection thresholds which are more sensitive to compounds found only in low levels.

We did not find a difference in the ability of males to produce pheromone compounds when reared with and without pollen in this experiment. This finding was somewhat unexpected, as pollen is an important resource for adult *Heliconius*. This result could mean that pheromones do not represent a condition-dependent signal. However, *Heliconius* are one of the most long-lived butterflies, with adults known to live more than 8 months in the wild (Gilbert, 1972) and pollen limitation might play a more important role over longer time scales. In females it has been shown that the effects of pollen for oviposition and viability are evident after about a month (Dunlap-Pianka *et al*., 1977), suggesting that until that point, larval reserves are sufficient. This could be the same for males with regards to pheromone production, with pheromones only being affected later in life when new energy resources are needed.

Males transfer nutrients to females during mating, which reduces the females’ need to forage for pollen (Boggs & Gilbert, 1979; Boggs, 1981, 1990). The spermatophore is protein-rich, and so the amino acids obtained by pollen-feeding are probably needed to make new spermatophores after each mating (Cardoso & Silva, 2015), as supported by the fact that males with more lifetime matings collect more pollen overall (Boggs, 1990). It has been proposed that females may determine spermatophore quality using cues, such as pheromones, that indicate direct benefits for the female (Cardoso & Silva, 2015). Alternatively, females could benefit indirectly through a “good genes” mechanism (Andersson & Simmons, 2006), for example through inheritance of foraging ability (Karino *et al*., 2005). Further experiments will be required to determine if pheromones in older males can act as indicators for spermatophore production and male quality, but our experiments do not provide any support for this hypothesis.

Chemical profiles produced by adult butterflies reared on the three larval host plants are largely overlapping. *H. melpomene* is able to produce the majority of compounds found in both androconial and genital bouquets when reared on all three *Passiflora* species. However, we find less variation in genital compounds produced by individuals reared on *P. menispermifolia*, the preferred host plant of *H. melpomene rosina*, compared to *P. vitifolia* and *P. platyloba*, perhaps suggesting some level of chemical or digestive specialisation.

Despite overall similarity between butterflies reared on different host plants, there are differences in some specific wing and genital compounds. Over one quarter of androconial compounds are found in significantly different amounts between the three groups, along with almost nine percent of genital compounds. One third of these significant wing compounds are thought to come from plants, originating from the lignin and lignane forming phenylpropanoid pathway (Boerjan *et al*., 2003). The latter are oxidized aromatic compounds with an alkyl sidechain up to three carbons, such as syringaldehyde, 1-(3,5-dimethoxy-4- hydroxybenzyl)ethanone or ethyl 4-hydroxy-3,5-dimethoxybenzoate. Over half of the genital compounds, specifically sesquiterpenes, are also thought to originate from plant sources. This suggests that differences in plant biochemistry affect the chemicals released from both androconial and genital regions of the adult butterfly.

We do not know which components of the androconial bouquet are biologically important in *Heliconius melpomene*. It cannot be assumed that the most abundant compounds are necessarily the most important, as minor compounds can often play important roles in attraction (D’Alessandro *et al*., 2009). Furthermore, the response to pheromonal cues is blend-specific in other Lepidopteran systems (Yildizhan *et al*., 2009; Larsdotter-Mellström *et al*., 2016). It is therefore possible that, despite the overlap in overall chemical composition between host plant treatments, the compounds which are significantly different could drive a change in response of the receiver. This seems particularly likely for the androconial bouquet, where more than a quarter of compounds are found in different amounts between larval host plant treatments. However, it is important to note that direct tissue extraction of pheromones may not accurately reflect chemical amounts emitted by live butterflies (Visser *et al*., 2018). Electrophysiological and behavioural experiments will be required to determine if these differences are biologically relevant.

Our results might contribute to an understanding of why some populations of *H. melpomene* are host specialists when their larvae can successfully feed on a wider variety of host plants. *H. melpomene* larval growth rate is similar on different host plants under laboratory conditions (Smiley, 1978), but there are slight differences in survival in the wild, perhaps due to ant attendance or parasitism (Merrill *et al*., 2013). In particular *H. melpomene* fed on *P. vitifolia* show a somewhat lower survival rate when compared with the natural host plant *P. menispermifolia* (Merrill *et al*., 2013). If host specialisation is not due to physiological adaptation in the larvae, an alternative is that it could be explained by pheromone differences between males reared on different host plants as larvae. If female choice acts on diet-derived pheromones, this could drive host plant specialisation through maternal-effect genes for oviposition choice (Quental *et al*., 2007). If selection is acting on the adult rather than the larval stage, then there is no expectation for larval host plant specialization. Furthermore, we might expect to find geographic differences in chemistry as *Heliconius* butterflies use different host plants across their geographic range and can vary in their extent of host plant specialisation (Benson *et al*., 1975; Benson, 1978). Both genetic and ecological differences between populations could result in differences in chemical bouquets.

As well as differences within species, compounds differ between species (Estrada *et al*., 2011; Mérot *et al*., 2015; Mann *et al*., 2017). Wing compounds are important for species recognition and mate choice (Mérot *et al*., 2015; Darragh *et al*., 2017), suggesting a potential role in reproductive isolation. It has long been suggested that male pheromones play a role in pre-mating barriers in *Heliconius* butterflies. Male *H. timareta florencia* and *H. melpomene malleti*, closely related co-mimics, court female wing models of both species equally, but interspecific mating is rare (Giraldo *et al*., 2008; Sánchez *et al*., 2015; Mérot *et al*., 2017). Another species pair, *Heliconius erato* and *H. himera* differ in the relative levels of investment in brain structures that process different sensory information (Montgomery & Merrill, 2017), which could suggest differences between species in the relative importance of olfactory cues. To understand the ecology and evolution of male pheromones in *Heliconius*, we have to understand what factors, both genetic and environmental, affect their production. Future work will allow us to investigate the role of chemical signalling in reproductive isolation between species and the underlying genetic architecture of these traits.

## Acknowledgements

We thank our team at the insectaries in Panama including Oscar Paneso, Sylvia Fernanda Garza Reyes and Diana Abondano. We also acknowledge the support and advice of William Wcislo. KD was supported by a Natural Research Council Doctoral Training Partnership and a Smithsonian Tropical Research Institute Short Term Fellowship. KJRPB and CDJ were supported by a European Research Council grant number 339873 Speciation Genetics. WOM was supported by the Smithsonian Tropical Research Institute and NSF grant DEB 1257689.

